# Premature termination codon readthrough in *Drosophila* varies in a developmental and tissue-specific manner

**DOI:** 10.1101/863027

**Authors:** Yanan Chen, Tianhui Sun, Zhuo Bi, Jian-Quan Ni, Jose C. Pastor-Pareja, Babak Javid

## Abstract

Despite their essential function in terminating translation, readthrough of stop codons occurs more frequently than previously supposed. However, little is known about the regulation of stop codon readthrough by anatomical site and over the life cycle of animals. Here, we developed a set of reporters to measure readthrough in *Drosophila melanogaster*. A focused RNAi screen in whole animals identified *upf1* as a mediator of readthrough, suggesting that the stop codons in the reporters were recognized as premature termination codons (PTCs). We found readthrough rates of PTCs varied significantly throughout the life cycle of flies, being highest in older adult flies. Furthermore, readthrough rates varied dramatically by tissue and, intriguingly, were highest in fly brains, specifically neurons and not glia. This was not due to differences in reporter abundance or nonsense-mediated mRNA decay (NMD) surveillance between these tissues. Overall, our data reveal temporal and spatial variation of PTC-mediated readthrough in animals, and suggest that readthrough may be a potential rescue mechanism for PTC-harboring transcripts when the NMD surveillance pathway is inhibited.

## Introduction

Fidelity of protein biosynthesis, including termination of the nascent polypeptide chain, is critical for all cellular functions. In eukaryotes, translation termination is a highly conserved process, and ribosomes can terminate protein biosynthesis with remarkable fidelity when encountering a stop codon^1,2^. Sometimes, however, translation can continue through a stop codon and terminate at the next in-frame stop codon, a mechanism commonly known as stop codon readthrough^3^.

In RNA viruses, readthrough is extensively utilized to append an extension domain to a proportion of coat proteins^4^. However, readthrough also occurs in the synthesis of eukaryotic proteins. In yeast, [*PSI*^+^] cells which carry the prion form of eRF3 (Sup35p) show a reduced efficiency of translation termination and higher rates of stop codon readthrough^3^. The [*PSI*^+^] strains exhibit phenotypic diversity which is beneficial to adapt to fluctuating environments^5,6^. In *Drosophila*, the *headcase* (*hdc*) gene encodes two different proteins, one of them produced by translational readthrough, and the readthrough product is necessary for *Drosophila* tracheal development^7^. There have also been a number of reports of readthrough in mammals, including in mouse brains^8-10^.

Validation of eukaryotic readthrough candidates had been confined to relatively small numbers until a comparative genomics methodology was used to analyze nucleotide sequences immediately adjacent to protein-coding regions in 12 *Drosophila* species. By identifying highly conserved sequences following native stop codons, Kellis and colleagues proposed more than 300 novel readthrough candidates^11^. Using ribosome profiling, Dunn *et al* experimentally validated a large number of these evolutionarily conserved readthrough candidates, as well as identifying more than 300 examples of non-conserved stop codon readthrough events in *D. melanogaster* embryos and the S2 cell-line^12^. Although there is some debate about whether stop codon readthrough truly represents a regulatory mechanism^13^, and there are mechanisms to mitigate canonical readthrough^14^, these data suggest that stop codon readthrough in eukaryotes is far more pervasive than previously appreciated. We therefore decided to measure readthrough in flies using a set of novel gain-of-function reporter lines that could sensitively detect translation through stop codons in animals throughout their life cycle, as well as in specific tissues. Furthermore, we confirmed that the stop codons in our readthrough reporters are recognized as premature termination codons (PTCs) in flies. We observed that stop codon readthrough frequency in two candidate gene reporters varied widely throughout fly development, and appeared to be highest in *Drosophila* neurons. High frequency readthrough of PTCs may be an alternative rescue pathway for translation of transcripts with premature termination codons in flies.

## Results and Discussion

### An *in vivo* gain-of-function reporter fly line can sensitively detect translational readthrough

We wished to measure stop codon readthrough in flies across developmental stages of their life cycle. We initially chose to measure readthrough using a candidate gene, *rab6*, which had been identified as undergoing moderately elevated readthrough rates in a ribosome profiling study of fly embryos^12^. We decided to use a gain-of-function reporter^15,16^, since this approach would be able to detect low levels of readthrough. The gene for Nanoluc luciferase – Nluc –^17^, missing its start codon, was cloned immediately downstream of *rab6* complete with its native stop codon and 3’ UTR, but missing the second, in-frame, stop codon (Fig. 1a). In our reporter, translation past the native UAG stop codon would result in functional Nanoluc luciferase enzyme, which could be detected using commercially available reagents (Fig. 1b). We were able to identify by tandem mass spectrometry a peptide derived from Nluc in flies expressing the reporter (Fig. 1c, d). To further confirm that readthrough was occurring, we raised a polyclonal antibody to a peptide coded by the 3’UTR of *rab6*, and verified that a protein band of the appropriate size was identified by Western blot (Fig. 1e). To verify Nluc protein was actually the product of translation past the stop codon, not alternative initiation of *nluc* translation bypassing the stop codon, we performed an “in-gel” Nanoluc luciferase assay capable of detecting functional enzyme on an SDS-PAGE separated whole-cell lysate (see Methods). We could detect functional Nluc as the major band corresponding to the size of the Rab6-translated-3’UTR-Nluc gene product, comparable to a control Rab6-Nluc fusion protein where the native TAG stop codon of *rab6* had been replaced by CAG, corresponding to a glutamine residue (Fig. 1f). Although some smaller bands were visible, they comprised a minority of the total signal, and may have represented alternate initiation or breakdown products. Taken together, these data confirm that Nluc was expressed in reporter flies as a result of stop codon readthrough.

**Fig. 1:**
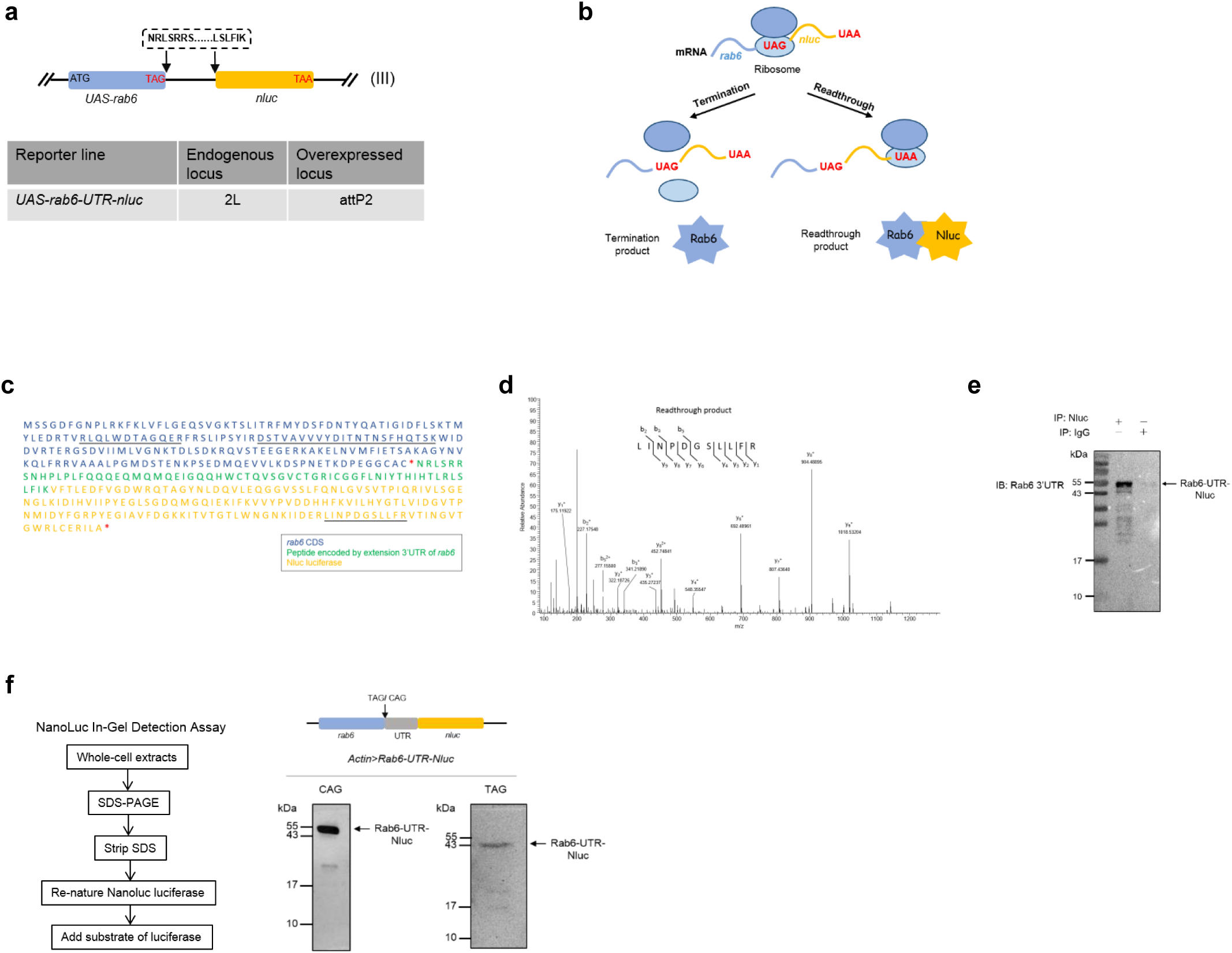
An *in vivo* gain-of-function reporter fly line can sensitively detect translational readthrough. **a**, Schematic for construction of the in-frame stop codon readthrough reporter. The gene coding for *nluc*, missing its start codon, was cloned immediately upstream of the second in-frame stop codon following *rab6*, its native stop codon and 3’ UTR. **b**, Schematic for basis of the stop codon readthrough reporter. The expression of UAS-Rab6-UTR-Nluc was driven by Actin-Gal4. Normal termination of translation at the first in-frame stop codon would result in no expression of Nluc. Translational readthrough would result in expression of Rab6-UTR-Nluc, and detection of luciferase activity. **c**, Primary sequence of the C-terminal extended polypeptide that would result from translational readthrough of the reporter. Red stars represent the stop codons. Underlined sequences represent peptides detected by mass spectrometry. **d**, MS/MS spectrum representing a peptide, LINPDGSLLFR, from Nluc. The spectrum contains a total of eight C-terminal “y” ions and three N-terminal “b” ions consistent with this sequence. **e**, Western blot with antibody raised against a 3’UTR peptide of Rab6 following IP of Nluc from the reporter fly line. **f**, Nanoluc luciferase in-gel detection identifies Nluc with migration consistent with translational readthrough. Whole-cell extracts of *Actin>Rab6-TAG-Nluc* adult flies and the variant with TAG- to-CAG substitution were assayed.

### The stop codon of reporter flies is recognized as a premature termination codon

We wished to use our reporter system to measure relative readthrough rates in a semi-quantitative way. To control for differential expression of the reporter, both the UAS-*rab6-UTR-* STOP-*nluc* reporter and UAS-*gfp* were driven by Gal4 (see Methods and Fig. S1a). Comparison of the GFP and Nluc ratios would allow for relative quantitation of stop codon readthrough^18,19^.

To investigate potential cellular mediators of readthrough in our reporter system, we crossed our reporter fly line with a focused sub-library of UAS-RNAi fly lines from the Harvard *Drosophila* RNAi Screening Centre (DRSC) Resource^20^ at Tsinghua. Knock-down was driven via an actin-promoter except where knockdown of the target gene via an actin-driver promoter was lethal, in which case knock-down was driven by *BM-40-SPARC-Gal4* and *Cg-Gal4*. We chose to define “hits” if Nluc/GFP signal was either two-fold decreased or increased compared with control. We screened a total of 615 candidate genes, and identified 90 potential regulators of *rab6* readthrough (Fig. S1b). To eliminate *rab6*-specific hits, we rescreened the hits with a second reporter line encoding a second readthrough candidate *rps20*^12^. Testing knock-down of these hits with the rps20 readthrough reporter identified 25 genes, which showed similar readthrough phenotypes when knocked down in both *rab6* and *rps20* lines (Fig. S1c-f and Dataset S1). Only one candidate, the nonsense-mediated mRNA decay (NMD) gene *upf1*^21,22^, showed increased Nluc/GFP in *rab6* and *rps20* reporters when knocked down (Fig. S2a).

Stop codons can be classified as ‘native’ or ‘premature’, and mechanisms of both termination and readthrough vary and depend on how the stop codon is recognized by the cellular machinery^22^. Nonsense-mediated mRNA decay, is an important surveillance mechanism for monitoring of transcripts encoding PTCs, and prevents their translation by rapidly degrading such transcripts. Our identification of *upf1* as a potential mediator of readthrough in our screen suggested that our reporter constructs were recognized as coding premature termination codons (PTCs). However, systematic investigation of Upf substrates in yeast and mammalian cells suggest that apparently normal mRNAs without the classical features of PTC-containing transcripts may also be targeted for degradation^23-25^. Was the native *rab6* transcript a non-canonical Upf substrate? Knock-down of *upf1* led to increased abundance of reporter mRNA (Fig. 2a), but the stability of native *rab6* mRNA was not significantly affected by *upf1* knockdown (Fig. 2b). These data suggested that rather than representing native gene readthrough, readthrough rates measured in our reporter flies represented variation in PTC suppression. To further confirm whether our reporters measured PTC readthrough, we knocked down two other NMD-associated factors, *smg5* and *upf3*^26^. Knockdown of both factors increased reporter abundance (Fig. 2c, d), as well as Nluc/ GFP measurements (Fig. S2b), verifying that the reporter constructs, but not the native genes are subject to NMD, and that relative readthrough rates measured using the reporters would represent PTC readthrough.

**Fig. 2:**
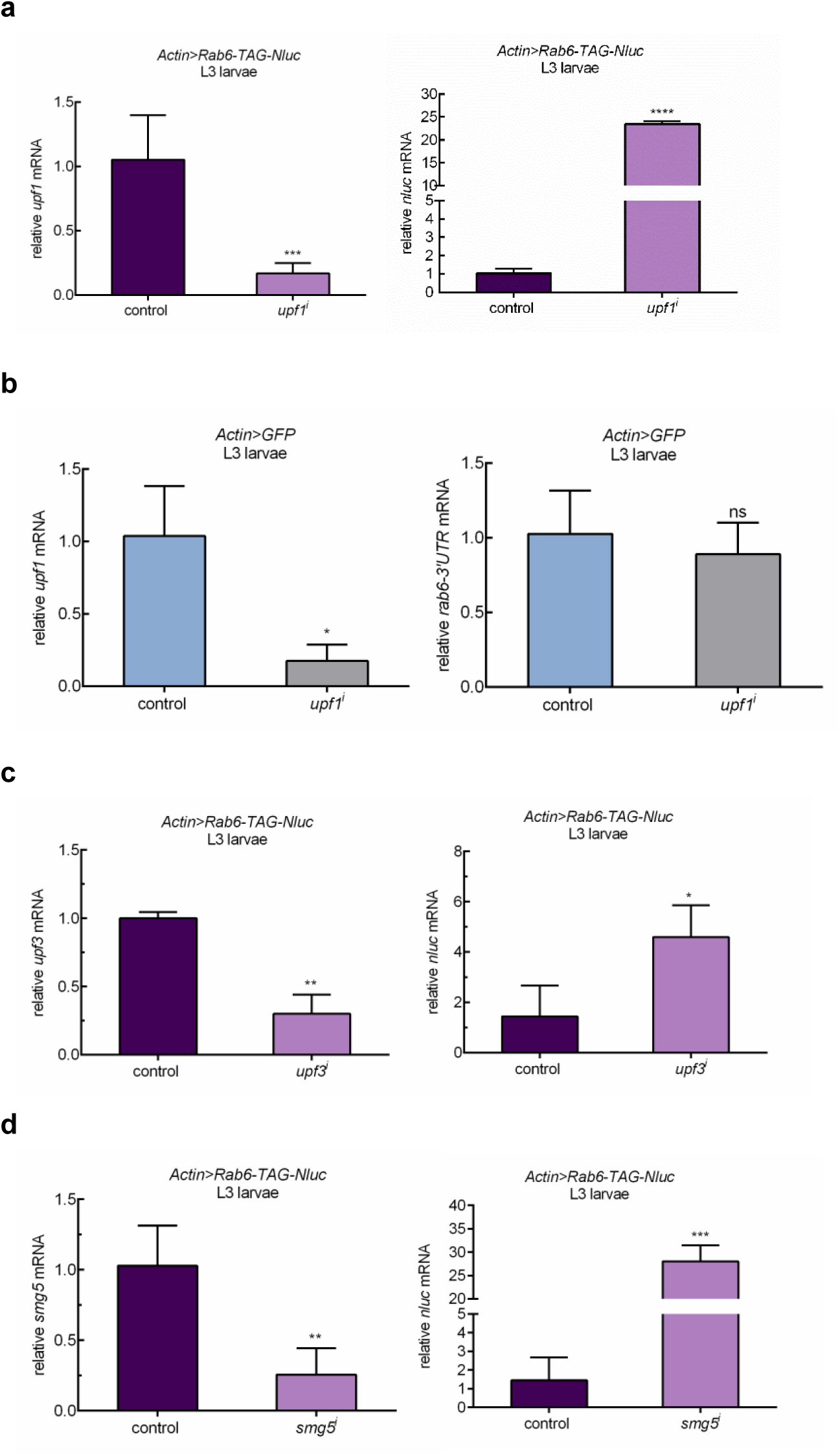
The stop codon of reporter flies is recognized as a premature termination codon. **a**, Knockdown of Upf1 is associated with increased abundance of *rab6*-based reporter transcript. *Actin>Rab6-TAG-3’UTR-Nluc* fly expressed both readthrough reporter and GFP. *upf1* (THFC_TH02846.N) was knocked down efficiently by Actin promoter (left panel). *rab6* reporter mRNA abundance was measured by quantitative RT-PCR with *nluc* primer. *rp49* abundance was used for normalization. **b**, Knockdown of Upf1 doesn’t affect abundance of endogenous *rab6. upf1* was knocked down efficiently by Actin promoter (left panel). Native *rab6* mRNA abundance was measured by quantitative RT-PCR with *rab6-3’UTR* primer. *rp49* abundance was used for normalization. **c, d**) Knockdown of NMD associated factors increases rab6-based reporter transcript. *upf3* (BDSC_58181) and *smg5* (BDSC_62261) were knocked down efficiently by Actin promoter. *rp49* abundance was used for normalization. Data represent means of three independent experiments ± s.d in a-d. \**P* < 0.05, \*\**P* < 0.01, \*\*\**P* < 0.001, \*\*\*\**P* < 0.0001; ns, *P* > 0.05 by Student’s t-test.

### PTC readthrough is regulated in a developmental stage-dependent manner

Since the initially-designed reporters were expressed as two constructs, and only one, expressing Nluc, but not the other expressing GFP was subject to NMD, it was possible that our apparent readthrough rates may be skewed, and that some of the hits from the RNAi screen represented interference with NMD and not readthrough *per se*. We therefore constructed a single-fusion reporter construct, representing *egfp-rab6-TAG-UTR-nluc* (Fig. 3a). Activity from the reporter was detected only when both Nluc and Gal4 were expressed (Fig. 3b), verifying its specificity and absence of background signal. The Nluc signal from whole-cell extract was derived almost entirely from a protein representing the correct molecular weight for the fusion product, which was strongly supportive that the reporter represented readthrough (Fig. 3c). Furthermore, immunoprecipitating and blotting against GFP identified two proteins with sizes representing normally the terminated translation, as well as a minority product (<1%) representing the extended polypeptide that would result from stop codon readthrough (Fig. 3d). To determine whether the new reporter was also subject to NMD, and therefore whether readthrough represented PTC suppression, we raised flies on cycloheximide or DMSO^27^. Cycloheximide treatment can stabilize mRNA transcripts subject to NMD^28,29^. Consistent with our other reporters, the new fusion reporter, but not native *rab6* also appeared to be subject to NMD (Fig. 3e-f). Having constructed a reporter that would measure relative PTC readthrough rates, we wanted to determine whether levels of readthrough in our PTC reporter varied by development stage (Fig. S3). We measured Nluc/GFP through larval development (Fig. 3g-h) and also when adults were hatched at day 0, through day 50 of adult life (Fig. 3i). In immature flies, readthrough increased, peaking at the pupa stage (Fig. 3h). In adult flies, readthrough increased from the newly hatched stage through to day 20, but decreased thereafter, although staying higher than the newly hatched adult (Fig. 3i), suggesting that increase in readthrough was not due to aging *per se*. These results were confirmed with a second fusion *egfp-rps20-TAA-UTR-nluc* reporter (Fig. S4).

**Fig. 3:**
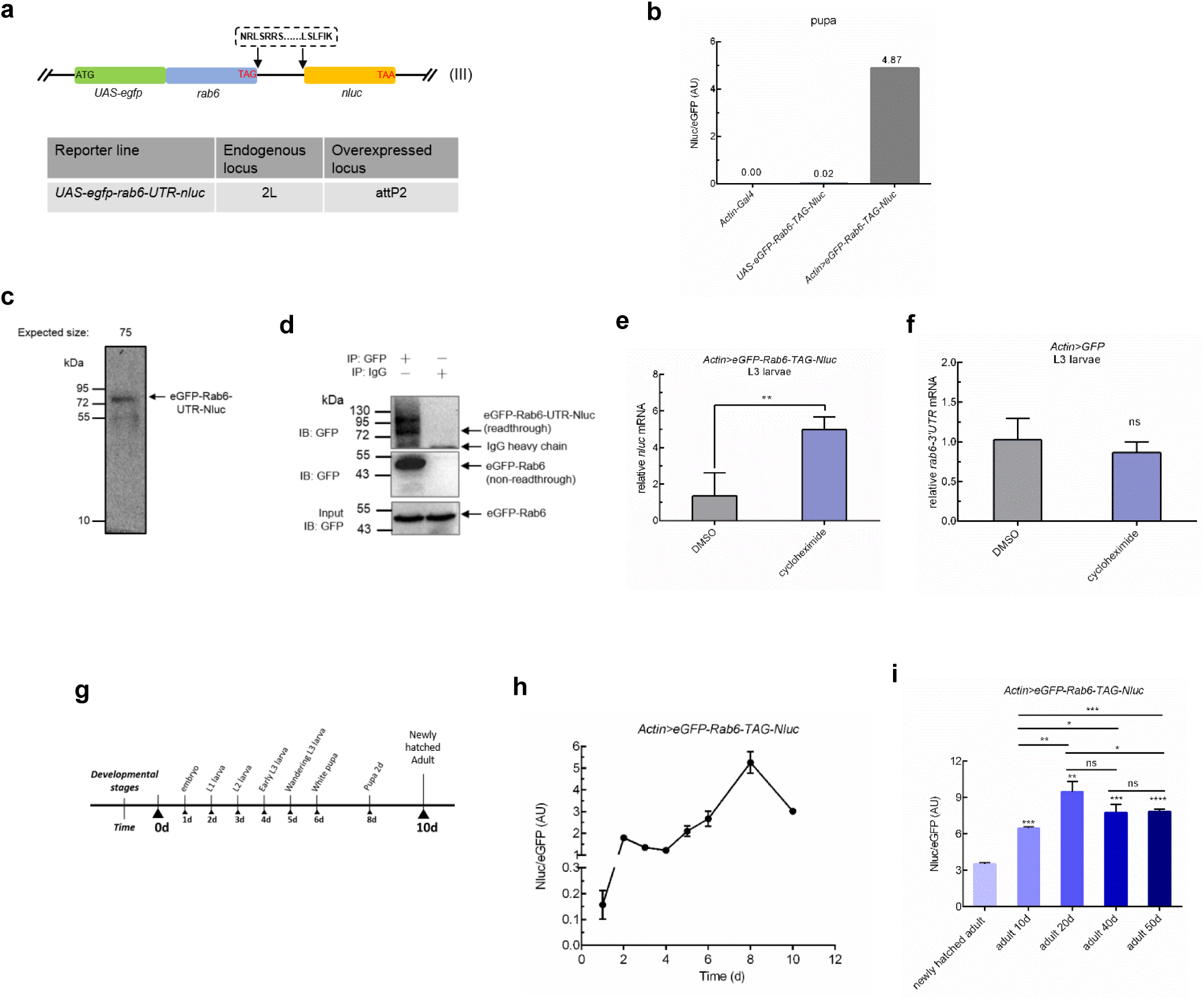
PTC readthrough varies in a developmental stage-dependent manner. **a**, Schematic for construction of the in-frame stop codon readthrough reporter. The gene coding for *nluc*, missing its start codon, was cloned immediately upstream of the second in-frame stop codon following *rab6*, its native stop codon and 3’UTR. Here, the peptide sequence NRLSRRS……LSLFIK were encoded by extension 3’UTR of *rab6*. To construct the transgene *egfp_rab6*_UTR_*nluc, egfp*, missing its stop codon, was fused immediately 5’ to *rab6* with its start codon removed. **b**, The readthrough reporter sensitively detects translational readthrough with minimal background signal. Nluc activity was only detected when the fly line expressed both the reporter and actin-driven Gal4. **c**, Readthrough product from *Actin>eGFP-Rab6-TAG-Nluc* whole-cell extracts was detected by Nanoluc luciferase in-gel detection assay. **d**, Readthrough efficiency was assayed by western blot. Lysates from *Actin>eGFP-Rab6-TAG-Nluc* adult flies were immunoprecipitated with mouse anti-GFP antibody or normal mouse IgG. eGFP-Rab6 (non-readthrough) and eGFP-Rab6-UTR-Nluc (readthrough) were separated by immunoblot analysis with rat anti-GFP antibody. PVDF membrane was cut at 55 kDa, eGFP-Rab6 was exposure with 10s and eGFP-Rab6-UTR-Nluc with 3600s, respectively. **e, f**, The reporter transcript of *rab6* is subjected to NMD. **E**, Suppression of NMD by cycloheximide increased readthrough reporter mRNA abundance. *Actin>eGFP-Rab6-TAG-Nluc* L3 larvae were transferred to standard fly food containing 500 μg/ml cycloheximide or DMSO for 24h. Reporter transcript abundance was assessed by real-time RT-PCR. **F**, Suppression of NMD does not affect endogenous *rab6*. Endogenous *rab6* mRNA abundance was measured by quantitative RT-PCR with *rab6-3’UTR* primer. *rp49* abundance was used for normalization. **g**, Cartoon representing time-points throughout the fly life-cycle when measurements were taken in (**h**). **i**, Higher readthrough level in old flies is not aging associated. Adult flies at day 10, 20, 40 and 50 of adult life were collected separately for analysis. Relative readthrough rates in *Actin>eGFP-Rab6-TAG-Nluc* adult flies were measured by normalized Nluc activity (Nluc/eGFP). Data represent means of three independent experiments ± s.d. \**P* < 0.05, \*\**P* < 0.01, \*\*\**P* < 0.001; \*\*\*\**P* < 0.0001; ns, *P* > 0.05 by Student’s t-test.

### Neuronal tissue undergoes higher rates of stop codon readthrough

To test the spatial variation of PTC-mediated readthrough, we constructed a series of reporter flies expressing the fusion reporters, driven by tissue-specific promoters. There were clear differences in levels of readthrough by larval tissue, with the highest PTC readthrough in brain tissue (Fig. 4a). To further confirm whether neuronal or glial cells were responsible for the high readthrough rates, we compared stop codon (PTC) readthrough in neurons with those in glia, by expression of the reporters in those tissues specifically. Neurons exhibited higher readthrough than glial cells (Fig. 4b, S5). Could differences in NMD and hence reporter mRNA stability explain this observation? In both neurons and glia, the reporter was subject was to NMD, but to a similar extent (Fig. 4c, d), and *nluc* transcript abundance was similar in both tissues (Fig. 4e), suggesting that neither tissue variation in NMD or transcript abundance were sufficient to explain the observed differences between glia and neurons. However, measured Nluc activity, corrected by GFP (Fig. 4b) or *nluc* transcript abundance (Fig. 4f) was higher in neurons than glia. Finally, we measured Nluc abundance, as well as the size of the Nluc-containing polypeptide by the Nluc in-gel assay, in reporters expressed in neurons or glia. For similar loading of either total protein or non-extended reporter (as measured by blotting against Actin or GFP respectively), there was substantially more Nluc, at a size corresponding to the readthrough product, in neurons compared with glia (Fig. 4g). These data confirm that PTC-readthrough in fly neurons occurred at higher rates than in fly glia.

**Fig. 4:**
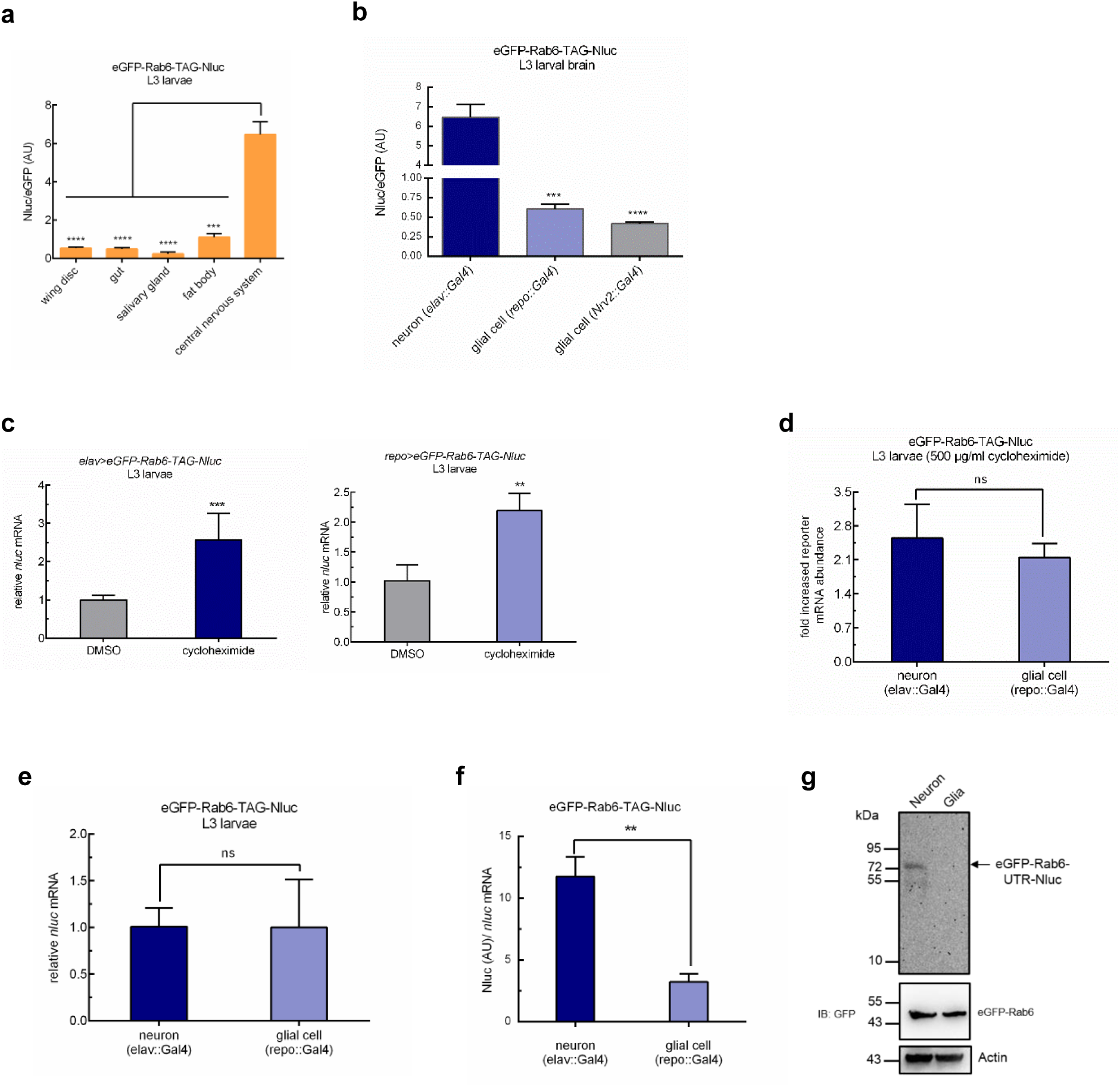
Neurons undergo higher rates of PTC readthrough. **a**, Rates of readthrough vary by tissue in wandering L3 larvae. Expression in wing disc was driven by *rn-Gal4*, gut by *crq-Gal4*, salivary gland by *He-Gal4*, fat body by *Cg-Gal4*, brain by *elav-Gal4*. Larval tissues were dissected to measure relative readthrough level. **b**, Neurons undergo higher rates of stop codon readthrough. Higher rates of translational readthrough in larval neurons compared with glial cells. Neuronal expression was driven by *elav-Gal4* and glial expression by *repo-Gal4* and *Nrv2-Gal4*, respectively. Larval brains were dissected and readthrough rates measured in cell-lysates. **c**, Reporter transcripts in larval neurons and glial cells are subject to NMD. L3 larvae were transferred to standard fly food containing 500 μg/ml cycloheximide or DMSO for 24h. Reporter transcript abundance was assessed by real-time RT-PCR. *rp49* abundance was used for normalization. **d**, The measured variation in Nluc/eGFP was not due to variation of NMD activity. Relative increase in transcript abundance in neurons and glia following suppression of NMD as measured by quantitative RT-PCR. *rp49* abundance was used for normalization. **e**, Reporter transcript abundance in neurons and glia. Reporter transcript abundance was assessed by real-time RT-PCR. *rp49* abundance was used for normalization. **f**, Neurons undergo higher rates of PTC readthrough. Relative readthrough level in neurons and glial cells in *eGFP*-*Rab6-TAG-Nluc* larvae. Nluc activity was normalized to the abundance of *nluc* mRNA, and *nluc* mRNA expression was determined by qRT-PCR. Data represent means of three independent experiments ± s.d. **g**, The measured variation in Nluc/eGFP are due to variation of readthrough rates. Neuronal expression was driven by *elav-Gal4* and glial expression by *repo-Gal4*. Readthrough products from *Neural cell>eGFP-Rab6-TAG-Nluc* whole-cell extracts were detected by Nanoluc luciferase in-gel detection assay. Loading control by Western blot. Data represent means of three independent experiments ± s.d. \*\**P* < 0.01, \*\*\**P* < 0.001, \*\*\*\**P* < 0.0001; ns, *P* > 0.05 by Student’s t-test.

## Discussion

### Higher readthrough rates in old flies is not associated with aging

Most studies on errors in gene translation have been on single-celled organisms, such as bacteria and yeast^18,30-33^ and therefore the relationship between translational error and development or anatomy has received limited attention. Mistranslation increases in response to stress, for example viral infection or oxidative damage^34^, and therefore it has been proposed that increased mistranslation over time may result in “error catastrophe” and may be one of the causes of aging^35-37^. Experimental evidence for this has been limited. Measuring fidelity of translation of polyU transcripts *in vitro* in brain homogenates from young and old rats, Filion and Laughrea failed to identify increased error rates with old age^38^. A more recent study found a strong positive correlation between fidelity of translation of the first and second codon and longevity in rodents^39^, suggesting that although mistranslation may not increase in old age, getting to old age may require high fidelity translation. However, of note, in that study, there was no correlation between stop codon readthrough and longevity^39^. In our study, whilst we found that older adult flies had higher rates of readthrough than newly hatched adults (Fig. 3i), this increase in readthrough did not increase further in very old flies (Fig. 3i), suggesting that some other regulatory factor than ‘old age’ may be responsible for the observation.

### Neurons are responsible for higher rate of PTC readthrough in CNS

Studies of tissue-specific translational regulation are still in their infancy^40^. A recent study measured the translatome by ribosome profiling of fly muscle through tissue-specific expression of tagged ribosomes^41^. Mutations in Rpl38 resulted in profound patterning defects due to tissue-specific expression of Rpl38 and its role in translation of Homeobox mRNAs^42^. CNS-specific expression of a mutated tRNA resulted in neurodegeneration in mice^43^, again verifying that components and regulation of the translation apparatus can be expressed in a tissue-specific manner. A recent study observed high rates of native stop codon readthrough in mouse brains, although differences in readthrough rates between neurons and glia were not apparent^10^. Our study suggests, in the context of premature stop codon readthrough, there may be substantial tissue-specific differences in mRNA translation. For both of our reporter gene constructs, there was increased readthrough specifically in neural tissue (Fig. 4). Further mechanistic studies are needed to explain the high neuronal readthrough in *Drosophila*.

### Readthrough may represent an alternative mechanism of PTC transcript rescue

Readthrough of the stop codon in our reporters mechanistically represented readthrough of a PTC (Fig. 2) and as such we cannot determine whether canonical readthrough rates also vary by developmental stage and anatomical site. In mammals, a stop codon is usually labelled as premature if it is located more than 50 nucleotides upstream of the last exon-exon junction^21,44^. However, in *Drosophila*, as well as yeast, the definition of a PTC occurs independently of exon-exon junctions^45^, and may include other elements for classification, such as “faux” 3’UTRs^46^. Long 3’ UTRs are permissive for targeting of non-PTC-containing transcripts as substrates of Upf1 and NMD^47-49^. It is therefore likely that by incorporating the *nluc* gene in the extended 3’UTRs of our reporter constructs, despite conserving all other elements of the stop codon context, the ‘native’ stop codons of *rab6* and *rps20* were re-classified as PTCs by the *Drosophila* cellular machinery.

Despite their potentially disastrous consequences and association with pathology^50^, premature termination codon-containing transcripts are surprisingly common. Alternative splicing of mRNA can result in a high frequency of PTC-containing transcripts^51^. And one study estimated that the typical human individual codes for approximately 100 nonsense- (PTC-containing) gene variants, of which 20 would be homozygous^52^. Since translation of proteins, abnormally truncated due to the presence of a premature termination codon could result in protein misfolding, all eukaryotic cells have evolved nonsense-mediated mRNA degradation as a quality control mechanism to target PTC-containing transcripts for destruction^22,44^. However, NMD is not 100% efficient, a significant proportion of PTC-containing transcripts escape NMD^53^. Remarkably, a recent comparison of homozygous PTC-containing genes in humans with their homozygous “wild-type” counterparts showed that mRNA and protein abundance were similar in the two groups, suggesting that escape of PTC-containing transcripts from NMD-mediated degradation may be the norm rather than the exception^54^.

The Upf-mediated NMD pathway has regulatory functions beyond surveillance of PTC-containing mRNA and may itself be repressed in a developmental manner^22,55^. Bioinformatic predictions have suggested that neural tissue may be particularly susceptible to mistranslation-misfolded protein-induced damage^56,57^. It was therefore surprising that a brain-specific microRNA, miR-128, actively repressed NMD^58^, and that this downregulation of NMD represented a switch that triggered neuronal development^59^. Potential mechanisms for rescue pathways for nonsense-containing transcripts that do not rely on NMD, include RNA editing^60^, alternative splicing, truncated proteins bypassing the stop codon, and readthrough of the stop codon^54^. Our data supports a model in which neuronal cells may still be protected from the potentially deleterious effects of translating PTC-containing transcripts, despite downregulation of NMD^61^ via enhanced readthrough of PTC-containing mRNAs.

## Materials and Methods

### Fly Genetics

Standard fly husbandry techniques and genetic methodologies were used to assess transgenes in the progeny of crosses, construct intermediate fly lines and obtain flies of the required genotypes for each experiment^62^. The Gal4-UAS binary expression system was used to drive transgene expression with temporal and spatial control^63^. Flies were cultured at 25°C, and anesthetized by CO_2_ prior to use in experiments. Fly strains used in this study are listed in Table S1. Intermediate strains were constructed using these strains.

**Table S1.** Fly strains used in this study.

| Fly strains | SOURCE | IDENTIFIER |
| --- | --- | --- |
| *D. melanogaster*: *w*<sup>1118</sup> | Bloomington *Drosophila* stock center | RRID: BDSC_3605 |
| *D. melanogaster*: *UAS-RpS20-TAA-Nluc (II)* | This study | N/A |
| *D. melanogaster*: *UAS-Rab6-TAG-Nluc (III)* | This study | N/A |
| *D. melanogaster*: *UAS-Rab6-CAG-Nluc (III)* | This study | N/A |
| *D. melanogaster*: *UAS-eGFP-Rab6-TAG-Nluc (III)* | This study | N/A |
| *D. melanogaster*: *UAS-eGFP-RpS20-TAA-Nluc (II)* | This study | N/A |
| *D. melanogaster*: *act-FO-Gal4 / TM6B* | Bloomington *Drosophila* stock center | RRID: BDSC_3954 |
| *D. melanogaster*: *actin-Gal4 / TM6B* | Gift from Jose | N/A |
| *D. melanogaster*: *Nrv2-Gal4* | TsingHua Fly Center | RRID: THFC_TB00131 |
| *D. melanogaster*: *elav-Gal4* | Gift from Yi Zhong | N/A |
| *D. melanogaster*: *repo-Gal4 / TM6B* | Gift from Yi Zhong | N/A |
| *D. melanogaster*: *crq-Gal4* | Bloomington *Drosophila* stock center | RRID: BDSC_25041 |
| *D. melanogaster*: *rn-Gal4 / TM6B* | Bloomington *Drosophila* stock center | RRID: BDSC_7405 |
| *D. melanogaster*: *BM-40-SPARC-Gal4 / TM6B* | Gift from Hugo Bellen | N/A |
| *D. melanogaster*: *Cg-Gal4* | Bloomington *Drosophila* stock center | RRID: BDSC_7011 |
| *D. melanogaster*: *He-Gal4* | Bloomington *Drosophila* stock center | RRID: BDSC_8699 |
| *D. melanogaster*: *UAS-Upf1.RNAi* | TsingHua Fly Center | RRID: THFC_TH02846.N |
| *D. melanogaster*: *UAS-Upf1.RNAi* | Bloomington *Drosophila* stock center | RRID: BDSC_43144 |
| *D. melanogaster*: *UAS-Upf3.RNAi* | Bloomington *Drosophila* stock center | RRID: BDSC_58181 |
| *D. melanogaster*: *UAS-Smg5.RNAi* | Bloomington *Drosophila* stock center | RRID: BDSC_62261 |

### Generation of Nluc-tagged readthrough reporter transgenic lines

To obtain *UAS-RpS20-TAA-Nluc, UAS-Rab6-TAG-Nluc* and *UAS-Rab6-CAG-Nluc* lines, the relevant *rps20-TAA-nluc, rab6-TAG-nluc* and *rab6-CAG-nluc* were separately cloned into vector pVALIUM10-roe using Gateway recombination. RNA extracted from S2 cells (a kind gift from Dr. Gong Cheng) was used to synthesize cDNA (Bio-Rad, cat#1708890). Coding sequences of *rps20, rab6* and their following extension *3’UTR*^12^ were PCR-amplified from cDNA template with primers adding *attB* site at the 5’ termini of the ORF. In order to increase the expression of the transgenes in *Drosophila*, a 5’UTR element *Syn21*^64^ and the Kozak sequence CAAA<u>ATG</u> (the start codon underlined)^65^ were added to the coding sequence. *nluc* was codon optimized by GENEWIZ (Nanjing), and both start codon and stop codon were removed, and this *nluc* gene was inserted before the second in-frame stop codon of the extension 3’UTR to acquire a fusion protein. Simultaneously, *attB* site was added to the 3’ termini of the modified fusion protein for subsquent Gateway cloning. Additionally, *rab6-CAG-nluc* was constructed by site directed mutagenesis using *rab6-TAG-nluc* as template. To construct *UAS-eGFP-Rab6-TAG-Nluc* and *UAS-eGFP-RpS20-TAA-Nluc, egfp*, missing its stop codon, was PCR-amplified and fused with *rab6-TAG-nluc* and *rps20-TAA-nluc*, respectively by overlap PCR.

The PCR products were purified by gel extraction (cwbiotech, cat#CW2302M) and recombined into vector pDONR221 (Life Technologies, cat#12536017) using Gateway BP Clonase (Life Technologies, cat#11789020). Then the entry clones were recombined with destination vector pVALIUM10-roe using Gateway LR clonase (Life Technologies, cat#12538120). The final plasmids UAS-RpS20-TAA-Nluc, UAS-Rab6-TAG-Nluc, UAS-eGFP-Rab6-TAG-Nluc, UAS-eGFP-RpS20-TAA-Nluc and UAS-Rab6-CAG were sent to Tsinghua Fly Center to obtain transgenic fly lines^20^ by site-directed insertion. To overexpress protein on the second chromosome, entry clone was integrated into attP40 loci, while the overexpressed protein on the third chromosome was obtained by integrating into attP2 loci^66^. All the primers used in transgenic fly lines construction are listed in Table S2.

**Table S2.**
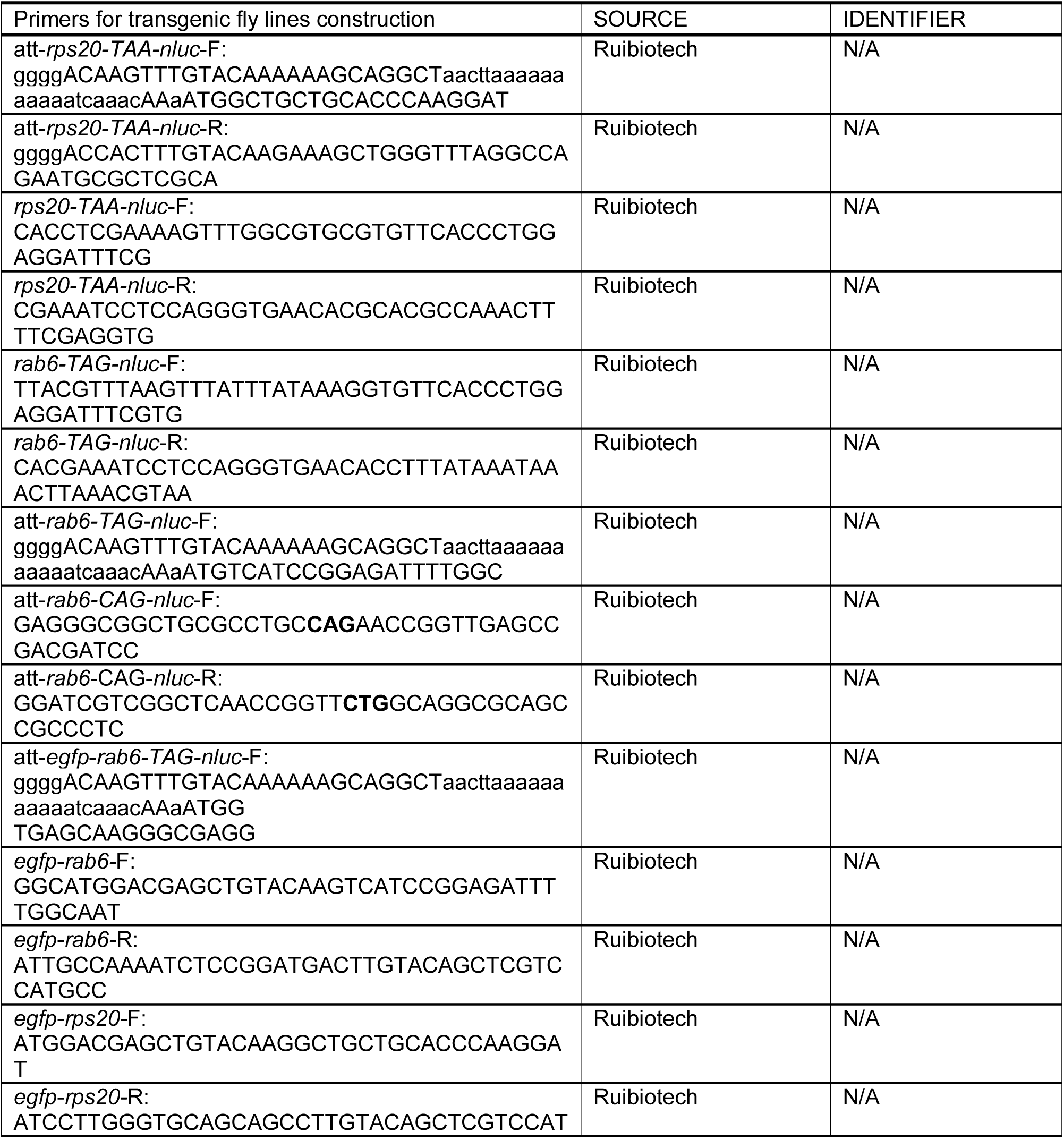
Primers used in transgenic fly lines construction.

### Luciferase-GFP assays

Luciferase was measured using the NanoGlo Luciferase Assay Kit (Promega, cat#N1120). Euthanized flies were collected in 200 μl of NB buffer (150 mM NaCl, 50 mM Tris-HCl pH 7.5, 2 mM EDTA, 0.1% NP-40) with addition of protease inhibitor cocktail (Biotool, cat#B14003), and homogenized with a 96-well plate multiple homogenizer (Burkard Scientific, BAMH-96). Homogenized samples were centrifuged at 20,000 rcf (4 °C) to pellet the larval remains. For measuring readthrough level (Nluc/GFP), 30 μl of each sample supernatant was transferred to a white-walled 96-well plate (Costar, cat#3922), an equal volume of Promega Luciferase Reagent was added to each well and incubated in the dark for 5 min. Another 30 μl of each sample supernatant was correspondingly transferred to a black-walled 96-well plate (Corning, cat#3925), and an equal volume of NB buffer was added to each well. Luminescence and fluorescence signal were measured by Fluoroskan Ascent FL (Thermo Scientific).

### Immunoprecipitation of Nluc

For western blot (Fig 1E), Nanoluciferase was immunoprecipitated by addition of 12 μg of rabbit polyclonal anti-Nluc IgG (a generous gift from Lance Encell, Promega) or normal rabbit IgG as a negative control (Cell Signaling Technology, cat#2729) respectively, to concentrated fly lysates, and incubated overnight at 4 °C to form the immune complex. The immune complex was captured by Protein A/G Plus Agarose (Pierce, cat#26146) and eluted by heating the resin with SDS sample buffer.

For mass spectrometry (Fig 1d), readthrough product of *rab6* was enriched by a Pierce Direct IP Kit (Pierce, cat#26148). For each enrichment, 10 μg of rabbit polyclonal anti-Nluc IgG (Promega) was coupled to AminoLink Plus Coupling Resin, and concentrated cell lysate was incubated with the resin overnight at 4 °C to form the immune complex. The enriched readthrough product was eluted by a neutral pH elution buffer (Thermo Scientific, cat#21027).

### Immunoprecipitation of eGFP-tagged readthrough reporter

For western blot (Fig 3d), eGFP-tagged readthrough reporter was immunoprecipitated by addition of 10 μg of mouse monoclonal anti-GFP IgG (Roche, cat#11814460001) or normal mouse IgG as a negative control (Proteintech, cat#66360-3-Ig) respectively, to concentrated fly lysates, and incubated overnight at 4 °C to form the immune complex. The immune complex was captured by Protein A/G Plus Agarose (Pierce, cat#26146) and eluted by heating the resin with SDS sample buffer.

### NanoLuc In-Gel Detection Assay

The protocol from the Promega technical manual (https://www.promega.com/-/media/files/resources/protocols/technical-manuals/500/nano-glo-in-gel-detection-system-technical-manual.pdf?la=en) was followed. Briefly, cell lysates were resolved on 15% SDS-PAGE. The gel was extracted from its casing and transferred to an appropriate tray. SDS was stripped from the gel by washing three times with 25% isopropanol in water (20 min each). NanoLuc was re-natured with three water washes (20 min each). The gel was developed with Furimazime (Promega, cat#N1120) / TBST (50 mM Tris-HCl pH 7.6, 150 mM NaCl, 0.05% Tween 20, 25 uM Furimazine). The gel image was captured on a white reflective background by Chemidoc XRS+ (Bio-Rad).

### Western blot

Western bolt was performed using standard methods. Rabbit polyclonal antibody to *Drosophila* Rab6 3’UTR was acquired by immunizing the New Zealand rabbit with peptide NRLSRRSNHPLPLFC by GenScript company (Nanjing). Readthrough product of *rab6* (Figure 1e) was detected using rabbit anti-Rab6 3’UTR antibody (2 μg/ml, GenScript) and revealed with Clean-Blot IP Detection Reagent (Thermo Scientific, cat#21230) and ECL Western Blotting Substrate (Pierce, cat#32106). For eGFP-tagged reporter Western blots (Figure 3d), elution was detected using 1:2500 dilution of monoclonal rat anti-GFP antibody (chromotek, RRID: AB_10773374), followed by a 1:5000 dilution of goat anti-rat IgG-HRP secondary antibody (easybio, cat#BE0109). Expression of reporter from input was detected by mouse anti-GFP IgG (Roche, cat#11814460001), followed by a 1:5000 dilution of goat anti-mouse IgG-HRP secondary antibody (cwbiotech, cat#CW0102). Loading control of neural cells (Figure 4g) was using 1:2000 dilution of anti-insect beta Actin mouse antibody (cmctag, cat#AT0008), followed by a 1:5000 dilution of goat anti-mouse IgG-HRP secondary antibody, or 1:2500 dilution of monoclonal rat anti-GFP antibody as mentioned before.

### Mass Spectrometry

Readthrough product of *rab6* was enriched by a developed IP method. After electrophoresis, SDS PAGE was stained by Imperial Protein Stain (Thermo Scientific, cat#24615), and the gel between 43 kDa to 55 kDa were cut, digested with trypsin, analyzed by Tsinghua Protein Chemistry Facility.

### Candidate forward genetic screen

Readthrough reporter fly line *Actin>Rab6-TAG-Nluc* or *Actin>RpS20-TAA-Nluc* was crossed with a set of *UAS-RNAi* fly lines (Tsinghua Fly Center) to knockdown target genes. Reporter fly line was crossed with *w1118*, and readthrough level (Nluc/ GFP) of progeny was as control. *BM-40-SPARC>RpS20-TAA-Nluc* and *Cg>Rab6-TAG-Nluc* reporter fly lines were utilized when knockdown target genes by *actin>Gal4* resulted in growth deficiency. For each knockdown genotype, 8-12 wandering L3 larvae were collected to measure readthrough level. Readthrough level of control was normalized to 1, and for each knockdown genotype, relative readthrough fold to control was calculated.

### Developmental stage assay

Parent transgenic reporter line *y v sc; UAS-RpS20-TAA-Nluc* (II) or *y v sc; UAS-Rab6-TAG-Nluc* (III) and an actin driven Gal4 line *w; Sp / CyO; act-FO-Gal4 UAS-GFP / TM6B* was crossed to express the readthrough reporter ubiquitously in the whole body of progeny. To eliminate the effect by variation in NMD efficiency, reporter line *y v sc; UAS-eGFP-RpS20-TAA-Nluc* (II) or *y v sc; UAS-eGFP-Rab6-TAG-Nluc* (III) was crossed with *y w; Adv / CyO; actin-Gal4 / TM6B*. After 6 hours of crossing, parent flies were removed to ensure the majority of the progenies were in the same developmental stage. Cell lysates were flash frozen by liquid nitrogen and stored at -80°C until all the samples of different developmental stages had been collected. Readthrough value was measured in whole cell lysate as above.

For embryo collection, standard apple juice agar plates were supplemented with fresh baker yeast paste, and collection cages were placed on the plates. Parent flies lay eggs on the plates for 4 hours. 0-4 hr embryos were collected from the agar plate using a small paintbrush.

For aged flies collection, newly hatched male and female flies were sorted into independent groups and cultured them in standard environmental conditions with a 12:12 hr light dark cycle. During the experimental period, flies were transferred to new vials containing fresh food every 2-3 days^67^. After 40 days of adult life, flies started to die and living flies would be transferred to fresh food every day to avoid the flies sticking to the food in the old vial.

### Protein extraction from *Drosophila* embryos

Collected embryos were washed gently in a collection basket (Corning, cat#352350). The base of the basket was dried by a paper tissue, and the basket was transferred to a container with 50% commercial bleach solution. Incubated for 5 min with gentle, periodic stirring to remove the chorionic membrane of the embryos. The dechorionated embryos became hydrophobic and floated on the surface of the bleach solution. Transferred the basket to a new container with deionized water, washed for 2 min and repeated twice. After wash, about 30 embryos of each independent sample were transferred to a 1.5 ml eppendorf tube containing 200 ul of ice-cold NB lysis buffer (protease inhibitor cocktail was added) and homogenized by a cordless motor (Kimble, cat#749540-0000). Homogenized samples were centrifuged at 20,000 rcf (4°C) to pellet the larval remains, and supernatant avoiding the upper lipid layer was transferred to a new 1.5 ml eppendorf tube.

### Protein extraction from different *Drosophila* organs or tissues

Expression of readthrough reporter in wing disc was driven by *rn-Gal4*, gut by *crq-Gal4*, salivary gland by *He-Gal4*, fat body by *Cg-Gl4*, neuron by *elav-Gal4*, glial cells by *repo-Gal4* and *Nrv2-Gal4*. Different larval organs or tissues were dissected and collected in 100 μl of NB lysis buffer with addition of protease inhibitor cocktail, homogenized by a cordless motor. Homogenized samples were centrifuged at 20,000 rcf (4 °C) to pellet the larval remains, and supernatant avoiding the upper lipid layer was transferred to a new 1.5 ml eppendorf tube. The final cleared cell lysates were flash frozen by liquid nitrogen and stored at -80 °C until all the samples of different *Drosophila* tissues or organs had been collected.

### Antibiotics treatment

L3 larvae were transferred to standard fly food containing 500 μg/ml cycloheximide (MedChemExpress, cat#HY-12320) ^27^ or DMSO for 24h. Afterwards, total RNA were extracted for real-time RT-PCR.

### Real-time quantitative RT-PCR

Approximate 10-20 wandering L3 larvae were collected and RNA was extracted by TransZol Up (TransGen Biotech, cat#ET111-01). DNase I (TransGen Biotech, cat#GD201-01) was used to digest genomic DNA. cDNA was synthesized with iScript Reverse Transcription kit (Bio-Rad, cat#1708890) and iTaq™ Universal SYBR Green Supermix (Bio-Rad, cat#1725120) was used for quantitative PCR. Analysis was performed in a CFX96™ Real-Time PCR Detection System (Bio-Rad). *rp49* was used as a reference gene^68,69^. All PCR reactions were performed in biological triplicate. Primers used were:

*rp49*-For: 5’-GGCCCAAGATCGTGAAGAAG-3’;

*rp49*-Rev: 5’-ATTTGTGCGACAGCTTAGCATATC-3’;

*upf1*-For: 5’-ACTTCCGGTTCGCACATCAT-3’;

*upf1*-Rev: 5’-CTTCCACTGTTCCTGGTCCC-3’;

*upf3*-For: 5’-ATGCTCCCTTCCAGTGCTTC-3’;

*upf3*-Rev: 5’-CCGCTTGATGAACTCCTGGT-3’;

*smg5*-For: 5’-GCTTTTTGACTGGCTGCGAA-3’;

*smg5*-Rev: 5’-ACCAGAGAATCACGCACGTT-3’;

*nluc*-For: 5’-GATCATCCCCTACGAGGGCT-3’;

*nluc*-Rev: 5’-GTCGATCATGTTGGGGGTCA-3’;

*rab6*-3’UTR-For: 5’-ATCCAACCATCCTCTCCCCC-3’;

*rab6*-3’UTR-Rev: 5’-GCAGATCCGGCCAGTACATA-3’;

*rps20*-3’UTR-For: 5’-ATCATCGACTTGCACTCGCC-3’;

*rps20*-3’UTR-Rev: 5’-GCACGCCAAACTTTTCGAGG-3’.

### Quantitation and Statistical Analysis

Most of experiments were performed at least three times on separate days (that is, independent experiments). Statistical analysis and graphic representation were performed with Graphpad Prism software. Two-tailed non-parametric Mann-Whitney tests were used in Fig S2A (data did not pass normality tests), otherwise means between two sample sets were compared by unpaired, two-directional Student’s *t*-tests in the rest of comparisons. \**P* < 0.05, \*\**P* < 0.01, \*\*\**P* < 0.001, \*\*\*\**P* < 0.0001 were considered statistically significant results.

## Acknowledgements

We thank Dr.Yi Zhong (*elav-Gal4 and repo-Gal4*) for kindly sharing fly strains; Lance Encell, Paul Otto and Thomas Machleidt from Promega for generously providing the anti-Nluc antibody and the protocol for the NanoLuc luciferase In-Gel Detection Assay; and the Tsinghua Fly Center for constructing transgenic *in vivo* readthrough reporter fly lines.

## Author Contributions

BJ conceived the project. YNC and BJ designed research. YNC performed the vast majority of experiments with assistance from THS and ZB. JQN provided the platform for generation of fly lines. JCPP and BJ supervised research. YNC and BJ wrote the manuscript with input from JCPP and other authors.

