## Supplementary material for "Premature termination codon readthrough in *Drosophila* varies in a developmental and tissue-specific manner": Supp Figures

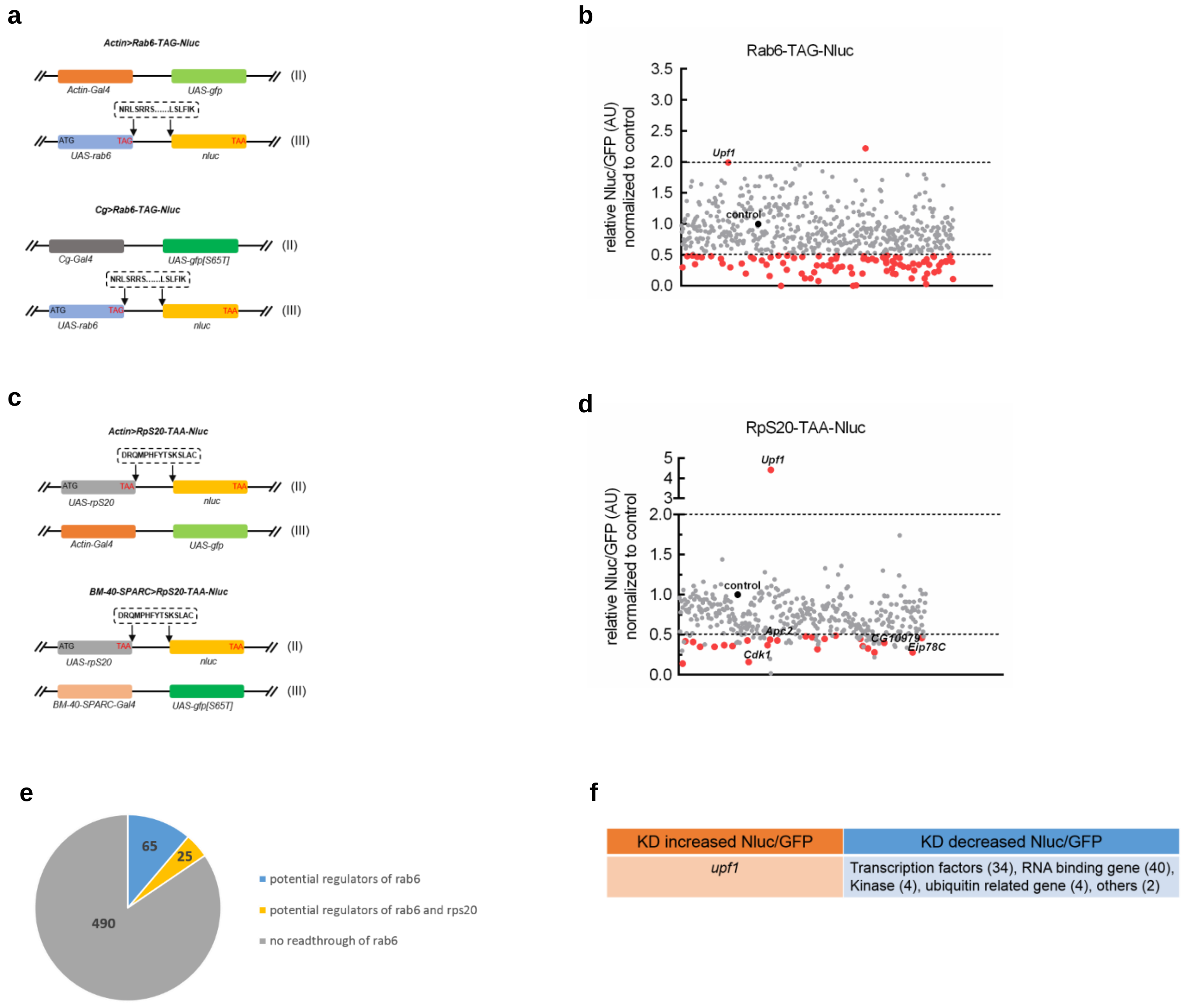


**Supplementary figure 1. A candidate forward genetic screen identifies potential regulators of *rab6* readthrough.**
**a**, Schematic of reporter fly line used in the primary forward genetic screen. The reporter fly line was crossed with a set of *UAS-RNAi* flies to knock-down target genes. **b**, Result of the candidate forward genetic screen identifying potential regulators of translational readthrough of *rab6*. The control (black dot) represents relative readthrough rate of the progeny of the reporter fly line (♀) and $w^{1118}$ (♂) and normalized to 1. Hits (above or below dotted lines) were defined as relative Nluc/GFP 2x higher or lower than control. **c**, Schematic of reporter fly line used in the secondary forward genetic screen. **d**, Result of the candidate forward genetic screen identifying potential regulators of translational readthrough of *rps20*. Hits of *rab6* were re-tested by rps20-based reporter, and red dots represent hits that showed the same result in both reporter fly lines. Results are summarised by the pie chart in **(e)**. **f)** *upf1* is implicated in increased Nluc/GFP signal in both reporters.

**a**

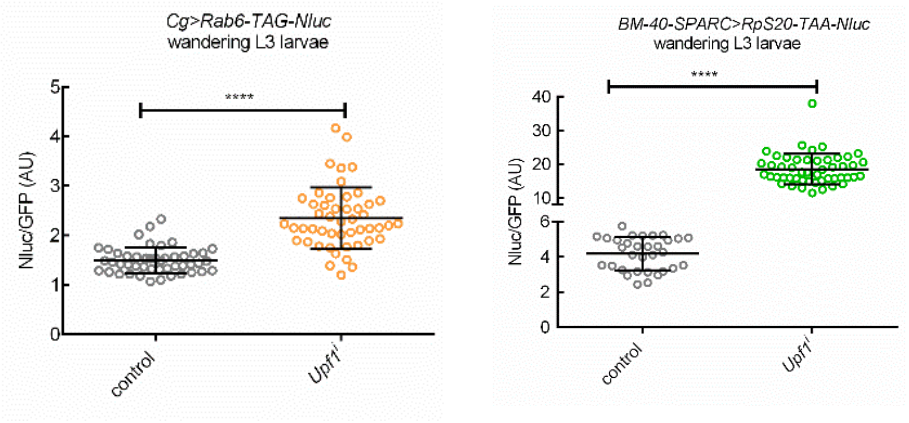


**b**

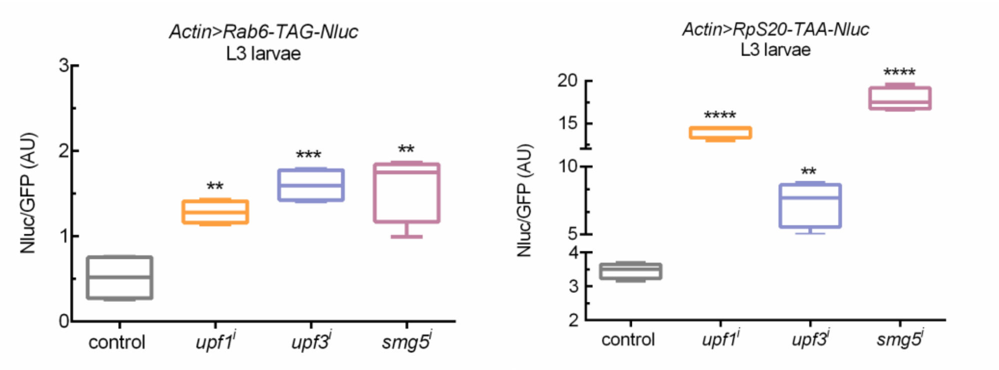


**Supplementary figure 2. Nluc/GFP reporters in flies are sensitive to NMD.**

**a**, Knock-down of *upf1* results in increased Nluc/GFP of both *rab6* and *rps20*. Each single wandering L3 larva was collected and readthrough level was independently calculated by measuring Nluc activity and dividing by GFP to control for variation in UAS-driven protein expression. Means between control and *upf1* knockdown groups were compared by two-tailed, non-parametric Mann-Whitney tests (data did not pass normality tests). **b**, Knock-down of NMD associated factors results in increased readthrough products of both *rab6* and *rps20*. *upf1* (BDSC_43144), *upf3* (BDSC_58181) and *smg5* (BDSC_62261) were knocked down efficiently by Actin promoter. Data represent means of three independent experiments ± s.d. in C. ***P* < 0.01, ****P* < 0.001, *****P* < 0.0001 by Student's t-test.

.

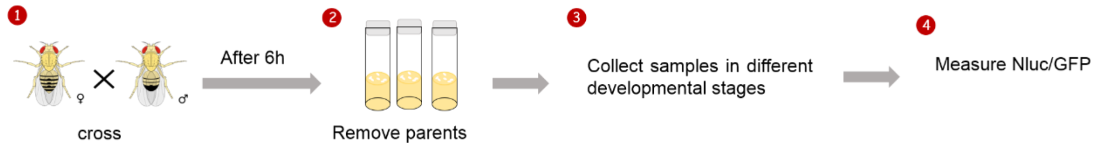


**Supplementary figure 3. Schematic showing how relative readthrough rates were measured over development stages.**
Parent transgenic fly lines *UAS-transgene-STOP-Nluc* and *Actin-Gal4* were crossed to generate the progeny *Actin>transgene-STOP-Nluc*. 6 hours following the cross, parents were removed to ensure progeny were all in a similar developmental stage. Readthrough rates were measured in cell-lysates of whole flies.

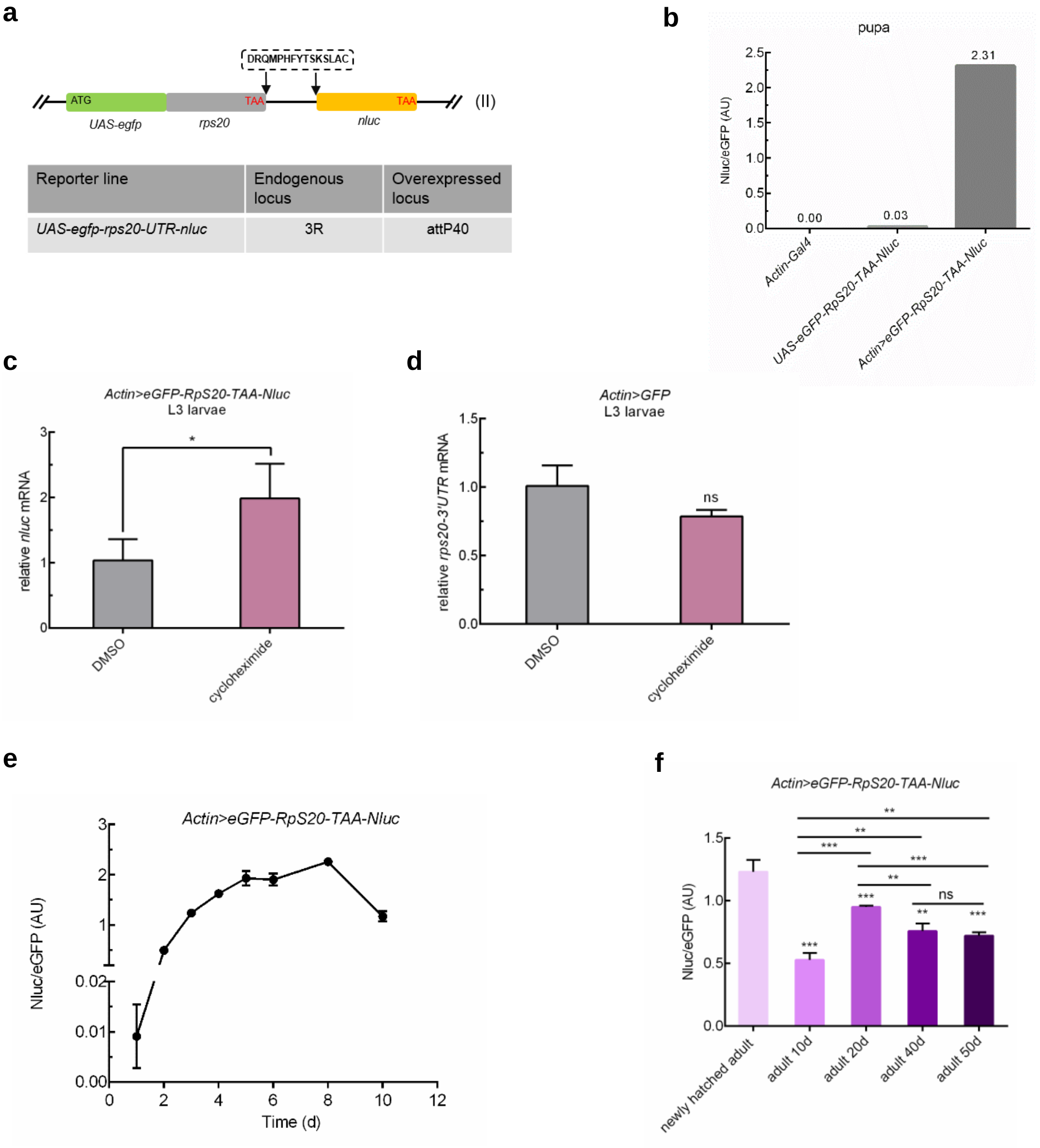


| Reporter line | Endogenous locus | Overexpressed locus |
|---|---|---|
| *UAS-egfp-rps20-UTR-nluc* | 3R | attP40 |

**Supplementary figure 4. PTC readthrough of *rps20* is also regulated in a developmental stage-dependent manner.**
**a**, Schematic for construction of the in-frame stop codon readthrough reporter. Here, the peptide sequence DRQMPHFYTSKSLAC were encoded by extension 3'UTR of *rps20*. **b**, As for the reporter measuring readthrough of *rab6* (see Figure 3B), Nluc was only detected when both the reporter and Gal4 were expressed. **c, d)** The reporter transcript of *rps20* is subjected to NMD. Reporter transcript abundance was assessed by real-time RT-PCR with *nluc* primer **(c)**. Endogenous *rps20* mRNA abundance was measured by real-time RT-PCR with *rps20-3'UTR* primer **(d)**. *rp49* abundance was used for normalization. **e**, Readthrough of *rps20* varied during fly life-cycle. **f**, Higher readthrough level in old flies is not aging associated.
Data represent means of three independent experiments ± s.d. **P* < 0.05, ***P* < 0.01, ****P* < 0.001; ns, *P* > 0.05 by Student's t-test.

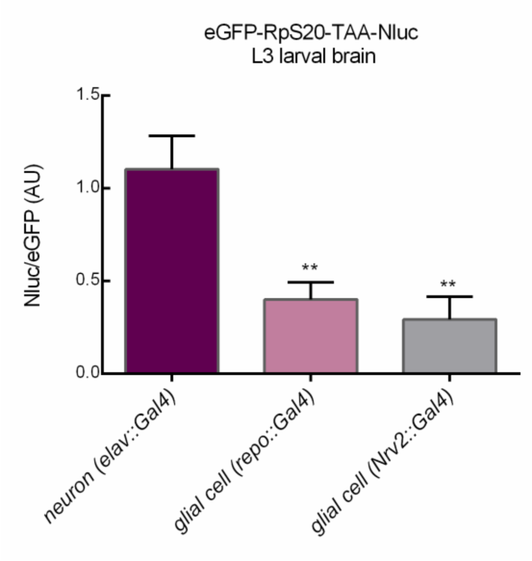


**Supplementary figure 5. PTC readthrough of *rps20* is higher in neurons**

*Drosophila* larval neurons undergo significantly higher readthrough level than glial cells. L3 larval brain were dissected. *rp49* abundance was used for normalization.

Data represent means of three independent experiments ± s.d. ***P* < 0.01 by Student's t-test.
